# Deep learning-based Drug discovery of Mac domain of SARS-CoV-2 (WT) Spike inhibitors: using experimental ACE2 Inhibition TR-FRET Assay Screening and Molecular Dynamic Simulations

**DOI:** 10.1101/2022.10.17.512637

**Authors:** Saleem Iqbal, Sheng-Xiang-Lin

## Abstract

SARS-CoV-2 exploits the homotrimer transmembrane Spike glycoproteins (S protein) during host cell invasion. Omicron, delta, and prototype SARS-CoV-2 receptor-binding domain show similar binding strength to hACE2 (Angiotensin-Converting Enzyme 2). Here we utilized multi-ligand virtual screening to identify small molecule inhibitors for their efficacy against SARS-CoV-2 virus using quantum Docking, pseudovirus ACE2 Inhibition TR-FRET Assay Screening, and Molecular Dynamic simulations (MDS). 350-thousand compounds were screened against the macrodomain of non-structural protein 3 of SARS-CoV-2. Using TR-FRET Assay, we filtered out two of 10 compounds that had no reported activity in *in-vitro* screen against Spike S1: ACE2 binding assay. Percentage Inhibition at 30 µM was found to be 79% for “Compound F1877-0839” and 69% for “Compound F0470-0003”. This first of its kind study identified “FILLY” pocket in macrodomains. Our 200 ns MDS revealed stable binding poses of both leads. They can be used for further development of preclinical candidates.

**Abstract Image:** 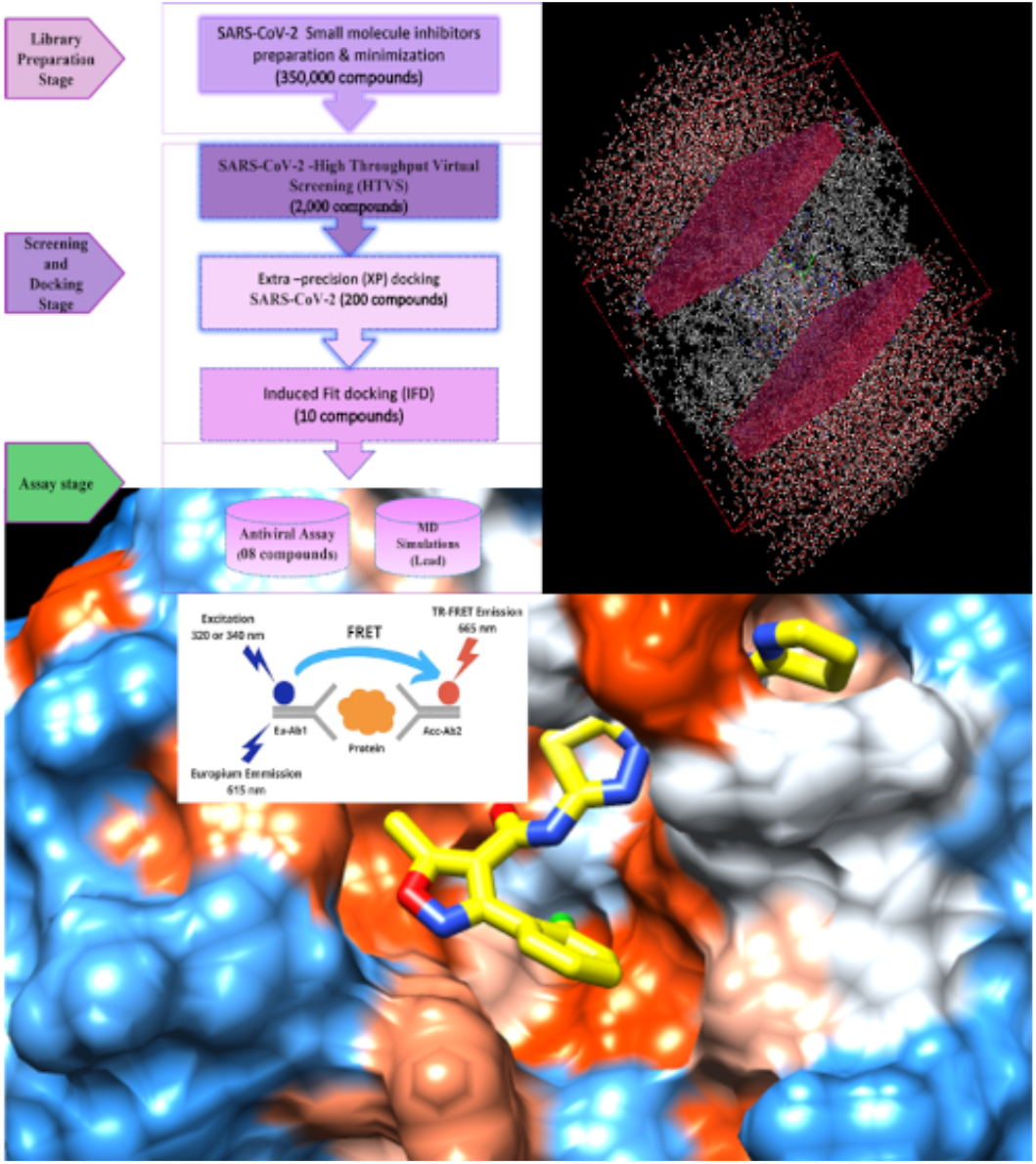

**In Brief:** Iqbal et al., described a deep learning guided drug discovery, efficacy against SARS-CoV-2 Spike inhibitors: using experimental pseudovirus ACE2 Inhibition TR-FRET Assay. Our molecular dynamic simulation results were next validated a posteriori against the corresponding experimental data of identified leads with 80 percent inhibition. Moreover, this study is first of kind to identify “FILLY” pocket in macrodomains.

**Highlights:**

- Experimental pseudovirus ACE2 Inhibition TR-FRET Assay and HTS lead to identification of two potential clinical leads.
- Conformational Dynamics analysis reveal the structural stability of complexes throughout 200 ns molecular dynamic simulations.
- Unveiling of the impact surface charge on the Variant of Concerns
- Detection of conformational changes within ACE2/RBD complex
- We identified the FILLY pocket in the SARS viruses.

## INTRODUCTION

The COVID-19 pandemic is a tremendous threat globally with many variants arising, some of which are variants of concern (VOC) including omicron (B.1.1.529), and its higher lineages. As of October 2022, 216 countries have reported COVID-19 cases, with more than 621 million confirmed and approximately 6,545,651 deaths (https://covid19.who.int/). More recently, Monkeypox virus and hemorrhagic fever virus (SHFV) which causes a lethal disease similar to Ebola virus disease has confronted global health emergencies too. Although several therapeutic agents have been evaluated for treatment, no efficacious antiviral agents have yet been shown. The causative agent of COVID-19, SARS-CoV-2 causes a lower respiratory tract infection that can progress to severe acute respiratory syndrome and even multiple organ failure (Meng et al., 2020; Yang et al., 2020). SARS-CoV-2 is an enveloped virus from the family Coronaviridae and genus beta-coronavirus, comprising a large positive-strand single-strand RNA (+ssRNA) genome (~30 kb), which encodes four structural proteins (spike, envelope, membrane, and nucleocapsid protein) that are components of the virus particle, 16 nonstructural proteins (Nsp) mostly with enzymatic activities and 6 accessory proteins (Yoshimoto et al., 2020; Finkel et al., 2020; Chan et al., 2020). Receptor binding is a key step of virus invasion (Cecon et al., 2022. Similar to severe acute respiratory syndrome coronavirus (SARS-CoV), SARS-CoV-2 uses its spike (S) protein to recognize the host receptor ACE2 (Lan et al., 2020). The SARS-CoV-2 utilizes ACE2 as the receptor for entry into target cells (Letko et al., 2020). Therefore, the S protein determines the infectivity of the virus and its transmissibility in the host (Hulswit et al., 2016). Studies carried out by (Han et al., 2022) has revealed that unlike alpha, beta, and gamma, omicron RBD binds to hACE2 at a similar affinity to that of the prototype RBD, which might be due to compensation of multiple mutations for both immune escape and transmissibility. Differential Sensitivity of the Natural Variants and Experimental Mutants to a Panel of Convalescent Serum Samples has been shown in Fig. S1(A).

SARS-CoV-2 keeps evolving into new variants due to sustained global transmission (Li et al., 2021a) (Refer Mutational landscape in Supplementary Section). Further, none of the variants and mutants demonstrated significantly altered sensitivity to all 10 convalescent sera, i.e., the EC50 values were not altered by more than 4-fold, irrespective of an increase or decrease, when compared with the reference strain (Fig. S1(B)). The point mutation-induced structural flexibility in S-protein, D614G, shifts the conformation of the S protein towards ACE2-binding fusion competent state and hence enhances SARS-CoV-2 infectivity in human lung cells (Yurkovetskiy et al., 2020). The SARS-CoV-2 macrodomain within the nonstructural protein 3 counteracts host-mediated antiviral adenosine diphosphate–ribosylation signaling. As the catalytic mutations render viruses nonpathogenic thus making this enzyme an antiviral target. Keeping such high data into consideration, with less impact of the mutation on RBD, we proceeded with nsp3 studies. COVID-19 can be controlled by designing small molecule drugs or monoclonal antibodies based on the process of viral binding to cell receptors. Due to the continuous emergence of new virus mutants, more drugs need to be screened.

The radial, rooted Phylogenetic Tree (PT) depicting the Genomic epidemiology of 2844 SARS-CoV-2 genomes sampled between Dec 2019 and May 2022 is shown in Fig. 1A, and Fig. 1B represents the time changes in the number of observations of SARS-CoV-2 throughout the world. Phylodynamics analysis of the viral genome as interpreted in Nextstrain presents a real-time view into the evolution and spread of a range of viral pathogens of high public health importance (Hadfield et al., 2014). PT depicting the Genomic epidemiology of SARS-CoV-2 represented by clock clade and rooted one have been displayed in Fig. 1C and Fig. 1D respectively.

**Fig. 1.**
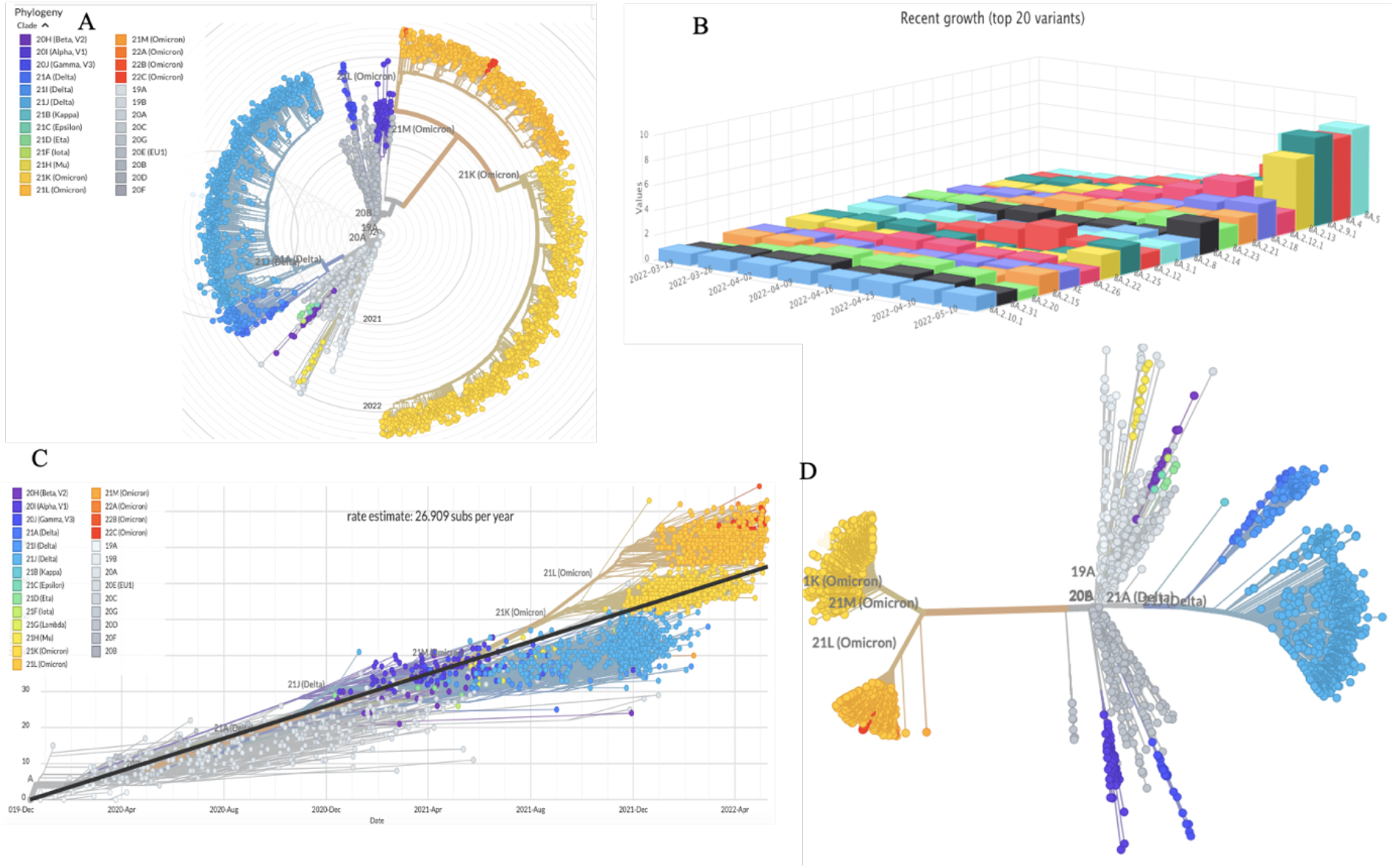
Illustration of genomic epidemiology of 2844 SARS-CoV-2 genome: Phylogenetic tree with Radial Clade depicting the SARS-CoV-2 Variants of Concern (VOC). (B) 3D representation depicting time changes in the no of observations of SARS-CoV-2 throughout the world. (C) Phylogenetic tree depicting the Genomic epidemiology of SARS-CoV-2 represented by clock clade. (D) Rooted phylogenetic tree depicting the Genomic epidemiology of SARS-CoV-2.

Nsp3 is the largest multidomain protein (~200kDa) in coronaviruses and is notable because of the presence of a key enzyme, papain-like cysteine protease (PLpro), which is essential for viral replication and a target protein for drug discovery (Imbert et al., 2008; Harcourt et al., 2004). The Nsp3 is found to be significantly different in two SARS-CoVs in comparison with other Nsps (Guo et al., 2020; Wu et al., 2019). The SARS-CoV-2 nsp3 includes three tandem macrodomains (Mac1, Mac2, and Mac3) (Srinivasan et al., 2020). The Genomic distribution of Missense and synonymous mutations of nsp3 is shown in the variation distribution plot in Fig. 2.

**Fig. 2.**
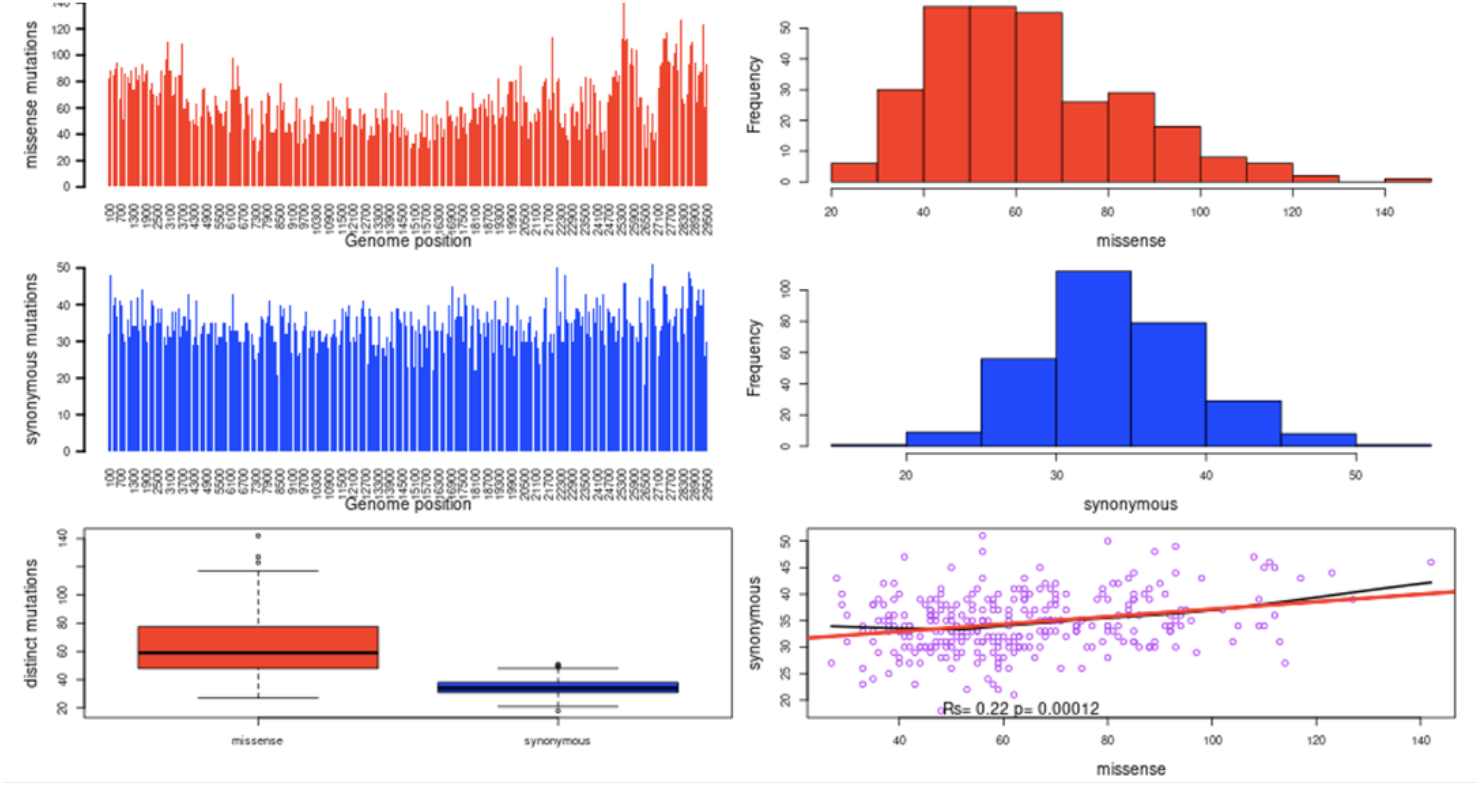
Variation Distribution plot of nsp3 depicting the Genomic distribution of Missense and synonymous mutations.

Mac1 is present in all CoVs, unlike Mac2 and Mac3, and early structural and biochemical data demonstrated that it contains a conserved three-layered α/β/α fold and binds to mono-ADP-ribose (MAR) and other related molecules (Alhammad et al., 2021; Egloff et al., 2006; Putics et al., 2005; Saikatendu et al., 2005; Cho et al., 2016; Xu et al., 2009). Macrodomains specifically recognize these modifications and serve therefore as “reader domains” of this posttranslational modification (Rack et al., 2016). Computational studies have shown that coronaviral Nsp3 may comprise 10-16 domains (Neuman et al., 2016; Lei et al., 2018). SARS-CoV-2 Nsp3 also includes multiple domains, an N-terminal ubiquitin-like (Ubl) domain followed by a highly variable and conserved macrodomain that binds to ADP-ribose (ADP) (Perina et al., 2014). It has recently been proposed that de-mono-ADP-ribosylation of STAT1 by ADRP may be linked to the Cytokine Storm Syndrome that is commonly observed in severe cases of COVID-19 (Claverie et al., 2020).

SARS-CoV-2 encodes the macrodomain (Mac1) domain within the large non-structural protein 3 (Nsp3), which has an ADP-ribosylhydrolase activity conserved in other coronaviruses. Mac1 shows highly conserved residues in the binding pocket for the mono and poly ADP-ribose. As revealed by Isothermal Calorimetry (ITC), Human CoVs bind to ADP-ribose with similar affinity (Fig. S2), where (A and B) depict the binding of human Mdo2 and SARS-Cov, MERS-CoV & SARS-CoV-2 Mac1 proteins, images in panel A correspond to two different experiments in ITC.

Therefore, currently SARS-CoV-2 Mac1 enzyme is considered an ideal drug target and inhibitors developed against them can possess a broad antiviral activity against CoV. Considering this, the ADP-Ribose-1’’-phosphate bound closed form of Mac1 domain is considered for screening with large commercial databases like Spec, Asinex, Life chemicals, ZINC, etc. We applied Extra Precision (XP) docking and Quantum polarized Ligand Docking (QPLD) providing strong potential lead compounds, that perfectly fits inside the binding pocket. QPLD algorithm begins with a Glide docking job that generates several geometrically unique protein-ligand complexes, Glide poses are subjected to charge refinement using QM/MM method (Schrodinger release 2021-4). A convolutional deep neural network-based approach called ‘Docking decoy selection with Voxel-based deep neural nEtwork’ - (DOVE) was used for selecting the protein model for docking (See Supplementary information). Trained on 2 million structures, DOVE considers atomic interaction types and their energetic contributions as input features applied to neural network. Using DOVE, we applied this convolutional neural network model to Crystal structure of SARS-CoV-2 spike receptor-binding domain with ACE2 (PDB entry 6M0J), SARS-CoV-2 macrodomain in complex with ADP ribose (PDB entry 6WOJ), (Table S1) and considering correct and acceptable according to CAPRI, a decoy of high probability (>0.5) and very small probability (<0.01).

Moreover, Laurini et al., 2020, performed computational-based simulations of the interaction between S-Protein region - receptor-binding domain (S-RBD) of SARS-CoV-2 and Angiotensin-Converting enzyme 2 (ACE2), highlighting residues playing an important role across human receptor/viral protein binding interface. During our research studies, parallel scientific study was published by (Han et al., 2022) reciprocating that Spike Omicron, delta, and prototype SARS-CoV-2 RBDs show similar binding strength to hACE2. A very recent study revealed that for neutralization of omicron, Monoclonal antibodies binding to receptor binding motif have been ineffective (Willett et al., 2022). Taking advantage of such an exemplary study, we proceeded with our screened candidates for checking effectiveness of compounds on WT Spike: ACE2 Binding. Thus, this manuscript focuses on parallel exploration of SARS-CoV-2 Spike inhibitors: using an experimental **Pseudovirus** ACE2 Inhibition TR-FRET Assay.

## RESULTS

### Binding of RBD of the SARS-CoV-2 spike protein to ACE2 monitored by TR-FRET Assay

Time-resolved FRET (TR-FRET) assays are increasingly used to monitor molecular interactions at nanometer scale with a high signal-to-noise ratio due to temporal separation between sample excitation and energy transfer measurements. TR-FRET combines proximity features of FRET assays with time-resolved fluorometry (Refer TR-FRET assay Supplementary Section). Pseudovirus ACE2 Inhibition TR-FRET Assay was carried out to check inhibitory effects of compounds against Spike: ACE2 binding (Refer to Supplementary information for experimental conditions (Table S3) and materials and methods). For list of ten TR-FRET assayed compounds and their test range used in this study, see WT Spike:ACE2 Binding assay (Table S2). Reactions were incubated for 1 hour at room temperature, and TR-FRET signal was finally measured in an Infinite M1000 microplate reader (Tecan) at excitation of 340 nm and emissions at 620 nm and 665 nm (Supplementary protocol).

### Inhibition of SARS-CoV-2 spike RBD binding to ACE2

Percentage inhibition is summarized in Table 1, Table 2. Moreover, Data for a reference inhibitor, Anti-Spike (0.0001 µM, 0.001 µM, 0.01 µM concentrations) was included as a control for inhibition. The values of percentage activity of compounds against Spike S1 (WT): ACE2 Binding were plotted on a bar graph (Fig. 3). “compound F2173-1125” was brightly coloured in the assay experiment and led to interference with TR-FRET signalling. At 30 µM concentration, percentage inhibition for all compounds revealed most potent inhibitor “F1877-0839” and showed 79 % inhibition which could be ascribed due to its possession of ribose scaffold moiety.

**Table 1.** Effects of Compounds on WT Spike: ACE2 Binding: Inhibitory effects of compounds against Spike: ACE2 binding are summarized in Table 1

| Inhibitor, Conc. | Percent Inhibition |
| --- | --- |
| F5084-0852, 30 μM | 0 |
| F1877-0839, 30 μM | 79 |
| F2619-0022, 30 μM | 30 |
| F1877-1292, 30 μM | 12 |
| F0466-0005, 30 μM | 0 |
| F2085-0027, 30 μM | 2 |
| F0772-2453, 30 μM | 43 |
| F0470-0003, 30 μM | 69 |
| F0827-0193, 30 μM | 38 |
| F2173-1125, 30 μM | 0 |
| Anti-Spike, 0.0001 μM | 21 |
| Anti-Spike, 0.001 μM | 49 |
| Anti-Spike, 0.01 μM | 76 |

**Table 2.** Data for the Effect of Compounds 1 - 10 on WT Spike: ACE2 Binding.

| Compound I.D. | TR-FRET Ratio |  | % Activity |  | % Inhibition |
| --- | --- | --- | --- | --- | --- |
|  | Repeat 1 | Repeat 2 | Repeat 1 | Repeat 2 |  |
| No Compound | 0.99 | 1.00 | 99 | 101 | 0 |
| F5084-0852, 30 μM | 1.00 | 1.03 | 103 | 111 | 0 |
| F1877-0839, 30 μM | 0.77 | 0.77 | 20 | 21 | 79 |
| F2619-0022, 30 μM | 0.89 | 0.93 | 62 | 77 | 30 |
| F1877-1292, 30 μM | 0.96 | 0.96 | 88 | 89 | 12 |
| F0466-0005, 30 μM | 1.01 | 0.99 | 104 | 97 | 0 |
| F2085-0027, 30 μM | 0.97 | 1.00 | 93 | 103 | 2 |
| F0772-2453, 30 μM | 0.89 | 0.86 | 62 | 51 | 43 |
| F0470-0003, 30 μM | 0.80 | 0.79 | 32 | 29 | 69 |
| F0827-0193, 30 μM | 0.89 | 0.89 | 63 | 61 | 38 |
| F2173-1125, 30 μM* | 1.15 | 1.20 | 153 | 172 | 0 |
| Anti-Spike, 0.0001 μM | 0.92 | 0.95 | 72 | 86 | 21 |
| Anti-Spike, 0.001 μM | 0.87 | 0.85 | 54 | 49 | 49 |
| Anti-Spike, 0.01 μM | 0.79 | 0.77 | 27 | 22 | 76 |
| Background | 0.71 | 0.71 |  |  |  |

\*Compound was brightly coloured in the assay, leading to interference with the TR-FRET signal in assay, leading to interference with TR-FRET signal.

**Fig. 3.**
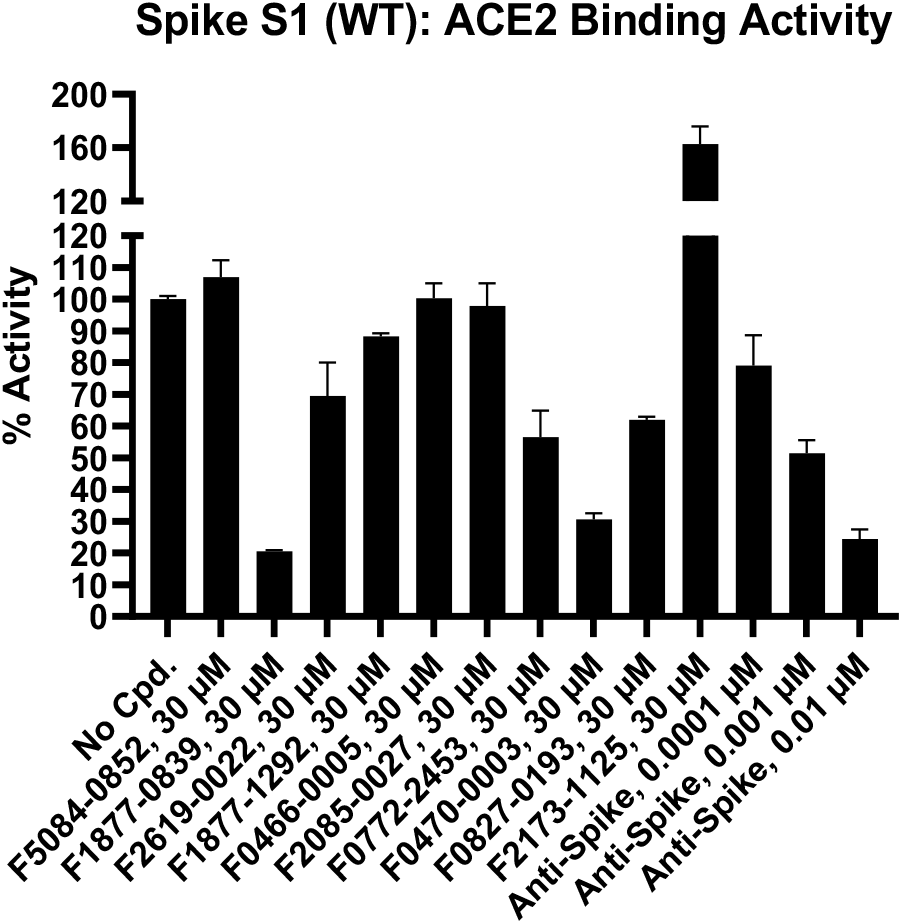
Percentage activity of compounds against Spike S1 (WT): ACE2 Binding

## DISCUSSION

Motivated by the fact that macrodomain can be inhibited by multiple drug-like ligands, and the drugs acting at protein-protein interfaces on spike protein (Kalathiya et al., 2020) and less-studied protein/domain viz., Mac1 domain of Nsp3 reducing viral entry to a mammalian host cell. We speculated that a range of drug molecules may efficaciously interact with ADP binding pocket. The Discovery and characterization of macrodomain inhibitors have been described here. To design ligands with high specificity and affinity it is crucial to understand structural determinants for protein-ligand complexes at an atomic level. We carried out all-atom MDS run for 200 ns and free energy calculations for top two hits that are widely used in studies of biomolecules, guided with fast and efficient deep learning (for selecting the target), and QPLD which provide change in angle conformations that bound more accurate inside binding pocket.

Besides identification of several high-potency drugs and/or molecules, we unveiled consensus binding mechanism that oxazole ring of both the drugs prefers to bind to core residue anchoring Phe 156, a residue novel to the SARS-Cov-2 virus family. Ligands anchoring at this site might facilitate future design and optimization of an inhibitor for the SARS-CoV-2.

In our modeling studies, to acquire comprehensive insight into putative inhibitor, a three-step docking procedure using Molecular docking filters such as (HTVS, XP docking) was used. After an in-depth analysis of binding patterns, third step in terms of induced fit docking (flexible) was employed. The glide docking of Schrödinger LLC Maestro package (New York, NY, USA) was initially carried out using Standard Precision (SP docking). Results were ranked by docking scores and were found to be −12.87, −9.61 & −11.19 kcal/mol for (F1877-0839) and (F0470-0003) & cocrystal-docked complexes respectively (Table S4). Inhibitory activity profound in compounds “F1877-0839” and “F0470-0003” may be attributed due to presence of a thiazole ring in both compounds. Moreover, both contain 1,3,4-thiadiazole ring at central core region of molecule and got suitably docked at distal side of binding pocket.

The docked-ligand molecule interaction within subsites has been shown in Fig. 4(A&B), ligand molecule interaction with the subsites. Adenine moiety is sandwiched between α2 and β7 in a mostly hydrophobic environment created by Ile23, Val49, Pro125, Val155, and Phe156. Polar contacts are facilitated by Asp22, which forms a hydrogen bond to N6 atom via its carboxylate group, and by main-chain amide of Ile23, which binds to N1 atom. Highlighting and striking insights in our study is that residue Phe156, which is replaced by Asn in closest homologues from SARS-CoV and MERS-CoV. Phe156 is one of important residues in SARS-Cov-2, and in our simulation studies lead inhibitors show consistent hydrogen bonding with such residue - Phe156 revealing the potency of the ligand-bound protein complexes of (F1877-0839) and (F0470-0003)(Fig. 4C&D). As depicted in Fig. 4 (E&F). The diphosphate moiety forms hydrogen bonds with key interacting residues Ser128 and Ile131. Phe132, Leu127, Ala129, Ala52, and Gly51 pave the way for hydrophobic interactions. Scaffold development of above-mentioned two inhibitors bound very well at the Mac1 active site providing critical structure-activity data for inhibitor optimization and our analysis reveals a path for accelerating the development of such inhibitors as potential candidates for antiviral therapeutics.

**Fig. 4.**
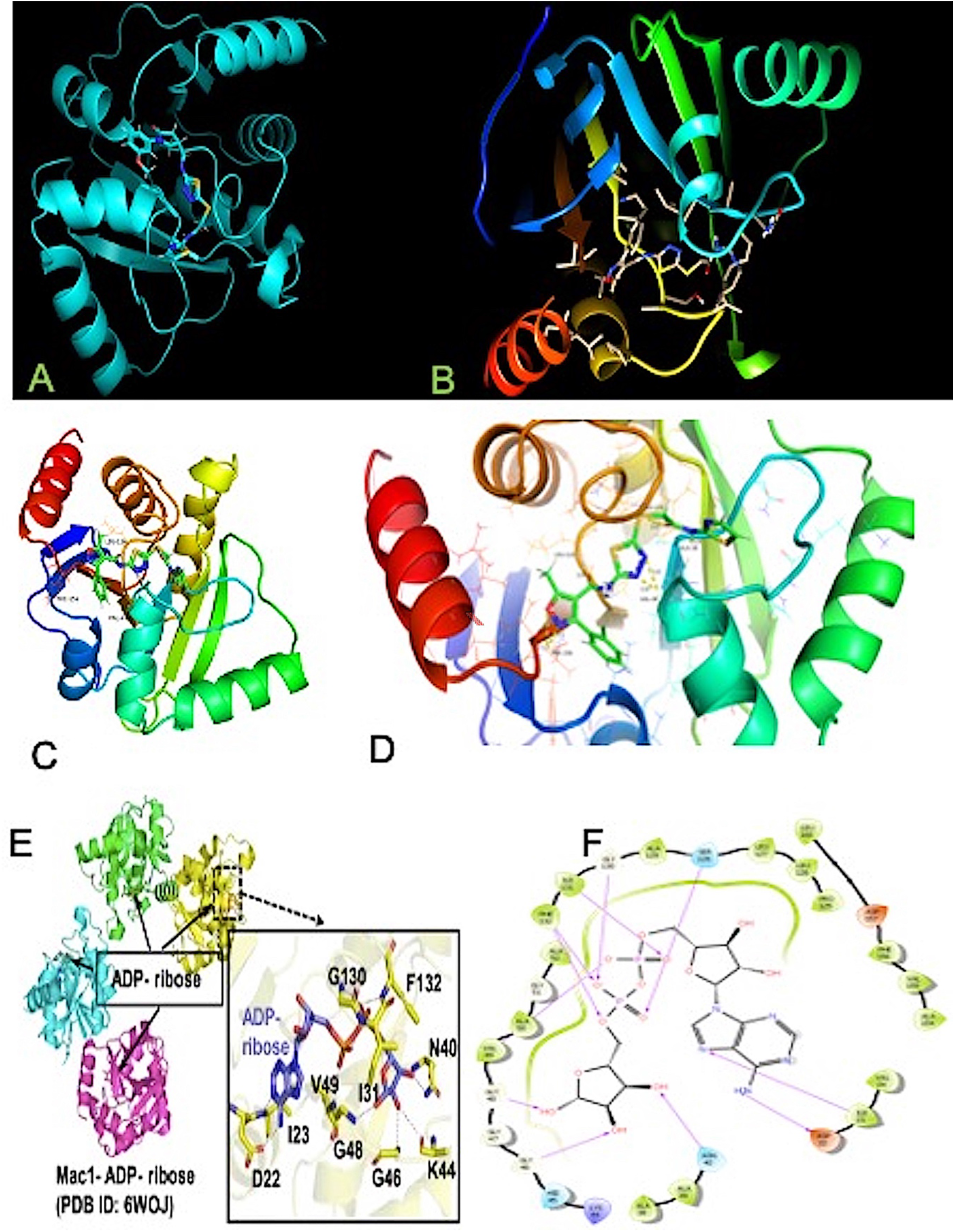
Docked complexes of “F1877-0839” & “F0470-0003” and their binding modes disclosed in the ribbon model. (A) cyan representation of docked complex of “F1877-0839” (B) docked complex of “F0470-0003”. (C) Hydrogen bonding interactions of compound “F1877-0839” at binding pocket of the macrodomain-ACE2 interface. (D) Zoomed in Catalytic site displaying key hydrogen bonding residues especially Phe156 and molecular interaction with subsites in the protein-ligand docked complex. (E) ADP-ribose bound at catalytic site while (F) represents the 2D-interaction showing various molecular interactions of the cocrystal.

As MDS complements experiments and provides structural information at atomic level with dynamics without facing the same experimental limitations. We first equilibrated crystal structure of SARS-CoV-2-ACE2 [Protein Data Bank (PDB) entry 6woj] in physiologically relevant environment. MDS was performed as previously described with slight modifications. MDS for 200 ns was run for final equilibrated structure (Supplementary Simulation methods). Further to understand stability of ligand in binding pocket of macrodomain, RMSD plots for the backbone atoms for both protein and ligand-bound protein complexes of (F1877-0839) and (F0470-0003). All runs for MD simulations of nsp3 macrodomain-ligand-bound complexes have been performed at the normal temperature. Structural deviations of each protein-ligand complex in terms of RMSD, RMSF for checking residual flexibility, and rGYR for compactness of each complex system, were investigated during MDS run of 200 ns. The stability of protein-ligand complexes was gauged during 200 ns. Fig. S4 represents the two-dimensional chemical structure representation of identified compounds used in study. For simulated docked complex containing compound (F0470-0003), average RMSD of protein Ca-backbone which was of magnitude of (0.6 A °), attained steady state after 100 ns simulation run while for compound (F1877-0839), remained steady for initial 50 ns with deviation of 1 A, and for rest 150 ns, showed average RMSD of 1.2 A °(Fig. S3). The average values for RMSF as well as rGYR are reported in (Fig. S3), and our overall simulation study reveal the stability of both the simulated complexes, signifying binding mode at the catalytic site of macrodomain.

Molecular interaction diagrams can be evidenced in terms of histograms and plots shown in Fig. 5, where residues Ala38, Gly48, Val49, Leu126, Ala129, Ile131, Phe132, and Phe156 contributed to hydrogen bonding interactions throughout the 200 ns simulation run. However, residues Val49, Phe132 and Phe156 interacting were more energetically favourable, consistent in terms of maintaining the hydrogen bond contact, and played important role in the regulation of binding the drugs at the catalytic site, paving the way for such class of compounds as potent inhibitors.

**Fig. 5.**
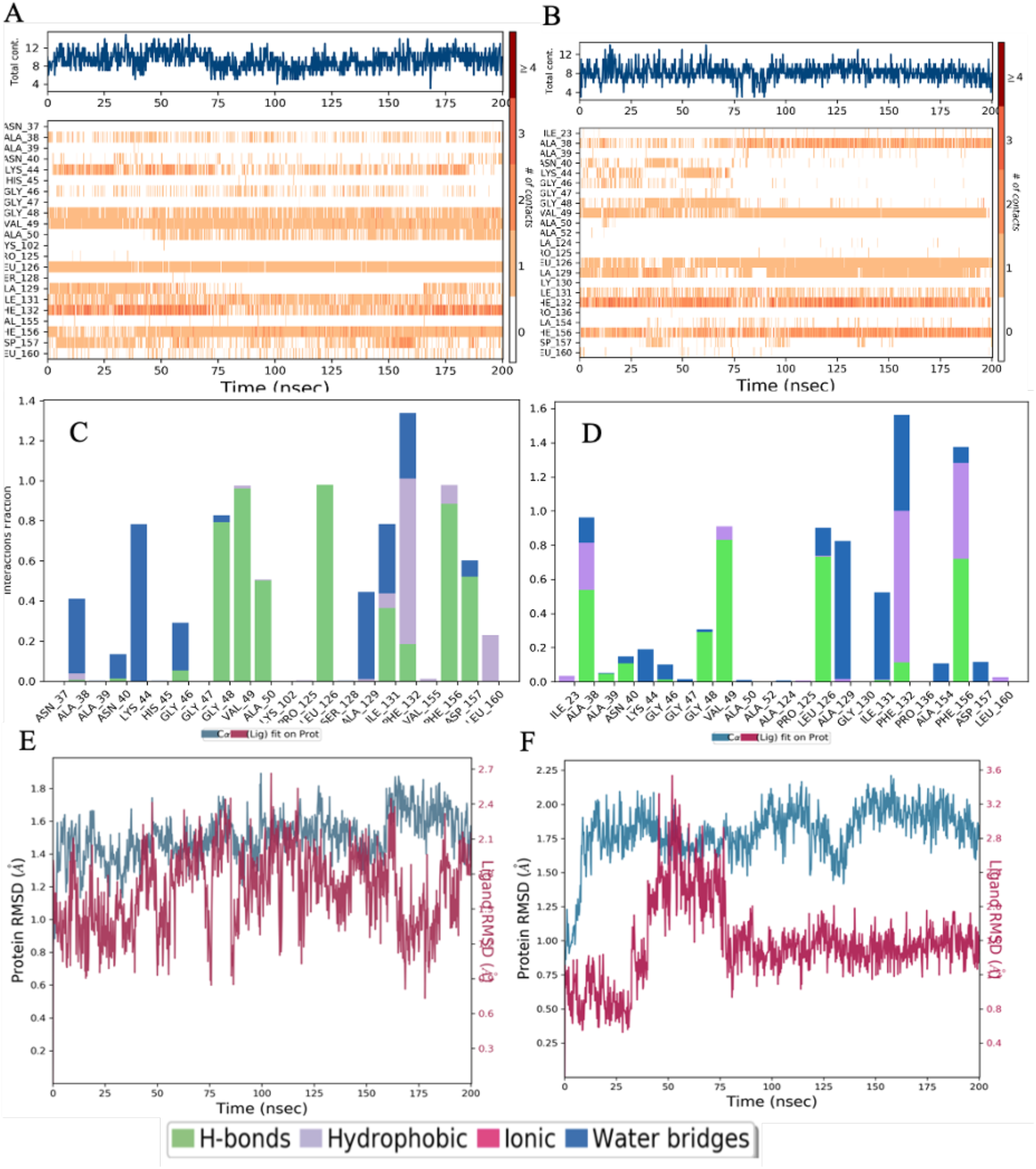
(A & B) Molecular interaction diagrams showing the consistency of hydrogen bonds of the key interacting residues. (C&D): Macrodomain (Mac) Catalytic site residues and active site hydrogen bond networks fractions of (F1877-0839) and (F0470-0003), (E&F): Corresponding RMSD’s of the protein Ca-backbone along with respective ligands F1877-0839 & F0470-0003.

Further to exemplify the conformational changes of every rotatable bond of both drug leads, torsional profile was calculated during simulation run of 0.00 to 200.00 ns. Fig. S5 (A&B) represents probability density of torsion of (F1877-0839) and (F0470-0003) illustrated in terms of dial plots, data so obtained has been plotted on bar plots (histograms) and provides insights into conformational strain ligand undergoes to maintain a protein-bound conformation (Fig. S5). Additionally, the protein RMSF, is useful for characterizing local changes along the protein chain has been shown (Fig. S5)(C&D).

To retain best compounds initially, Filters like Rule of Five, Drug Likeness, and ligand-based Absorption, Distribution, Metabolism, and Excretion/Toxicity prediction (ADMET) profiling aids in decreasing potential risks during clinical development (Table S5), and other molecular descriptors were taken into consideration. Interestingly, RMSD of the protein’s backbone calculated against the crystal (initial) one (PDB entry 6WOJ) saturated at 1.7 Å after ~10 ns. One caveat to our study is that we used monomeric forms of RBD and ACE2 for MDS which revealed that Spike protein was highly unstable. The native Spike protein is a trimer and has several other domains, including the nearby N-terminal domain, and the native ACE2 protein can exist as a dimer.

Conformational Dynamics analysis of the structural stability of complexes throughout 200 ns molecular dynamic simulations was estimated by calculating the protein-ligand RMSD. RMSD of the protein provides us insights into its structural conformation throughout the simulation. RMSD analysis discloses if the simulation has equilibrated. Both the simulated complexes were subjected to stability analysis. As shown in Fig. 5(E&F), displays corresponding RMSD’s of the protein Ca-backbone along with respective ligands F1877-0839 & F0470-0003 (Supplementary section for conformational Dynamics). This Molecular dynamic study further confirmed our TR-FRET assay where inhibition of the drug (F0470-0003) was less than the (F1877-0839), due to much conformational change in the aforementioned drug thus imparting its less stability than the potent drug-(F1877-0839) attaining high stability.

The use of electrostatic potential surfaces is a common approach for mapping complementary interaction interfaces in biomolecular complexes (Weiner et al., 1982; McCoy et al., 1997). We used structural modeling to demonstrate that macro domain is attracted to ACE2 receptors by long-range electrostatic forces leading to efficient recognition of the receptor (Fig. 6). Our results agree with recent studies by (Gan et al., 2022) for the gradual increase of +ve charge for emerging variants including higher lineages of omicron. The catalytic site of the macrodomain as per electrostatic calculations is concerned (Fig. 6) when the ligand binds when rotated to 180° the surface is red showing its high +ve charge. To our knowledge, our mapping identified a novel circular pocket, highlighted with the dark circle (Fig. 6) formed by residues Ser 5, Phe(F)6, Tyr(Y)8, Leu(L)11, Ile(I)17, Val29, leu136, Leu138, and Tyr 150, when rotated translationally. We coined the pocket with a single-letter amino acid name and designated it as the “FILLY” pocket. Residues surrounding the pocket are Arg 139 and Val 149. A short black arrow pointing toward the enclosed site in Fig. 6 discloses the identified pocket. Do such residues in the “FILLY” pocket represent any allosteric site or aid in glycosylation or represent a channel? needs scientifically investigated to ameliorate its biological significance.

**Fig. 6.**
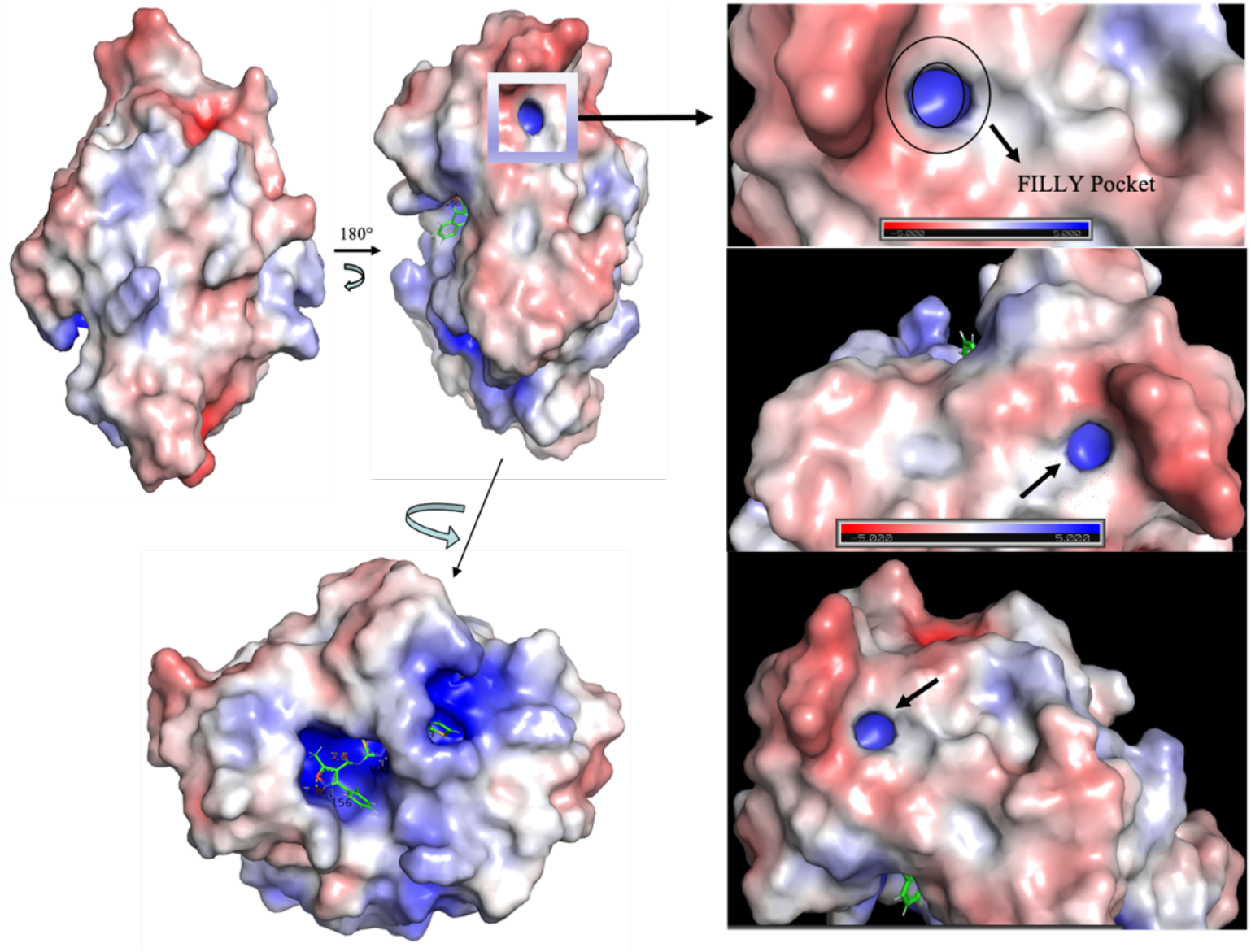
Electrostatic surface representation of the macrodomain bound compounds at its catalytic site. Where small single-direction arrows disclose the novel pocket; FILLY pocket.

Interestingly, as shown in the inset of Fig. 5(A-D), we found that during the equilibration, significant role of branched-chain amino acids such as Valine (Val), Leucine (Leu), Isoleucine (Ile), and aromatic amino acids Viz., Phenylalanine (Phe) whose levels are high in COVID-19 hospitalized patients, such residues got equilibrated during molecular dynamics simulations aiding us to reciprocate our identified leads to be potent inhibitors.

It is of great interest in modern drug design to accurately calculate the free energies of protein-ligand or nucleic acid-ligand binding. MM-PBSA (mol. mechanics Poisson-Boltzmann surface area) and MM-GBSA (mol. mechanics generalized Born surface area) have gained popularity in this field.

To calculate free energy and energy decomposition per residue and the Δ G_bind,solve_ MMGBSA calculations (Supplementary section for the methodology and the binding affinity). In Table 3, compound F1877-0839 showed higher binding free energy (ΔG _Total_ = −106.38 kcal/mol) than that of another compound viz., F0470-0003 and cocrystal, which showed binding affinity of −99.68 kcal/mol & −92.01 kcal/mol respectively. Better efficacy of combined strategy of molecular docking and free energy calculations to predict the binding-free energy indicates that F1877-0839 may be used as lead discovery and optimization of macrodomain targeting SARS-CoV-2.

**Table 3:** Free energy calculations (MM-GBSA) of identified lead compounds and Cocrystal.

| Name of compound | Solv GB | vdW | Coulomb | Covalent | Hbond | $\Delta G_{\text{Total}}$ (kcal/mol) |
| --- | --- | --- | --- | --- | --- | --- |
| F1877-0839 | 29.66 | -72.14 | -233.35 | 14.97 | -0.86 | **-106.38** |
| F0470-0003 | 48.16 | -54.17 | -40.41 | 1.23 | -0.78 | **-99.68** |
| Cocrystal | 23.75 | -62.55 | -31.65 | 6.7 | -7.78 | -92.01 |

Note: The bold values represent the best among the given factors.

## CATALYTIC MECHANISM

In order to decipher the structural information on the binding sites, it is important to know the Catalytic Mechanism involved at drug-protein interface **(See Supplementary section in detail for Catalytic mechanism involved**). For “Compound ID: F1877-0839” residues viz., Gly48, Val 49, Ala50, Ile131, Phe156, Asp157 contributed via hydrogen bonding while interacting residues such as Val49, Leu126, Phe156 interacted via hydrogen bonding in “Compound ID: F0470-0003”.We hypothesize it is the ribose scaffold of both compounds, induce the conformational change, compete with ADP-ribose binding site paving the way for key interactions with Phe156 and disclosed potency inhibition of 79% for “Compound ID: F1877-0839” and 69% for “Compound ID: F0470-0003”. Moreover, differences in both core structures lies in substitution of chlorobenzyl ring in “Compound ID: F0470-0003” while substitution of oxygen at ortho position and this varying electron density might impact long range electrostatic forces for receptor recognition. The peculiarity of our study identified water networks which play important role in protein ligand recognition, however their contribution to ligand binding is often difficult to identify. As shown in Fig. S3(C) and Fig. S3(D), we were able to identify the water networks. In continuation to such contribution’s residues viz., Ala38, Lys44, Ala129, Phe132 are involved in bridging interactions in “Compound ID: F1877-0839”. However, water mediated interactions are also profound in “Compound ID: F0470-0003” with Ala38, Ala129, Ile131, Phe132 residual contributions. Water networks reorganize upon ADPr binding, with a network of tightly bound water molecules acting as protein-ligand bridges, moreover tightly bound water molecules are co-opted by fragment binding, with implications for inhibitor design. Ligand-induced conformational changes in the SARS-CoV-2 ADRP structure was prevalent in the simulated complex.

**In conclusion**, present study illustrates molecular modelling insights into SARS-CoV-2 (WT) Spike inhibitors: using experimental ACE2 Inhibition TR-FRET Assay, high throughput Screening and MDS. MDS results validated a corresponding experimental data of identified leads and found Compound ID: “F1877-0839” as the best drug among all, inhibiting Spike protein in micromolar range, thereby making it therapeutic potential agent to treat Coronavirus. In parallel, this study is first of the kind to identify novel pockets in SARS-CoV-2 dubbed as FILLY pocket. Are FILLY pockets involved in transport pathway or exploring such passage to decipher the protein engineering functions, needs to be investigated. Further, our studies are under process of how changing the functional group of the screened compounds identified through pseudovirus inhibition assay, can impact the biological activity. Moreover, our findings could further be improved by exploring the other lead candidates by leveraging drug-target interactions along with experimentally known bindings by use of genetically expression datasets in addition to the Clinical Trail Data outcomes could be amicable for drug selection.

## SIGNIFICANCE

Covid-19 is triggered by infection with SARS-CoV-2 Virus, wherein Interaction between Receptor binding domain of the SARS-spike protein on surface of viral particle and the its receptor present on cell surface of human cells, the angiotensin I converting enzyme 2 (ACE2). Our findings suggest that electrostatic interactions are a major contributing factor for increased omicron transmissibility. Our structural modelling studies revealed that Spike receptor binding domain (S RBD) Plays pivotal role in enhancing ACE2 recognition. The peculiarity of our study lies in identification of water networks which play important role in protein ligand recognition. Our molecular dynamics simulation results validated a corresponding experimental data of identified leads and found Compound ID: “F1877-0839” as the best drug among all, inhibiting Spike protein in micromolar range, thereby making it therapeutic potential agent to treat Coronavirus. Moreover, this study is first of the kind to identify novel pockets-FILLY pocket in SARS-CoV-2.

## LIMITATIONS

With the unprecedented no of deaths as the experts forecast for next several years and the emergence of variants, the mechanisms of SARS-CoV-2 entry into host cells still remains mystery, and could be further explored by focussing on different targets for which our studies are under process. A more focus on exploring other receptors viz., Kidney Injury Molecule-1/T cell immunoglobulin mucin domain 1 (KIM-1/TIM-1), tyrosine-protein kinase receptor UFO (AXL), lectins, Cathepsins etc., will further help in exploring their importance which might help in deciphering their important role in promoting viral infection of the human respiratory system and indicate their role as alternative receptors for future clinical intervention strategies.

## Abbreviations

MDS: Molecular Dynamics Simulations
ACE2: Angiotensin-Converting Enzyme 2
TR-FRET: Time Resolved Forster/Fluorescence energy transfer
HTVS: High-Throughput Virtual Screening
XP: Extra Precision
QPLD: Quantum Polarised Ligand Docking

## Notes

### Competing Interest Statement

The authors have declared no competing interest.

