## Supplementary Info for "Deep learning-based Drug discovery of Mac domain of SARS-CoV-2 (WT) Spike inhibitors: using experimental ACE2 Inhibition TR-FRET Assay Screening and Molecular Dynamic Simulations"

*** Corresponding Address:**

1. **Prof. Sheng-Xiang-Lin**
   Full Professor and Senior Scientist
   Axe Molecular Endocrinology and Nephrology,
   CHU Research Center and Laval University,
   Quebec City,
   Quebec, G1V 4G2, Canada
2. **Dr. Saleem Iqbal**
   **Senior Postdoctoral Fellow**
   Department of Molecular Medicine,
   CHU Research Center and Laval University,
   Quebec City,
   Quebec, G1V 4G2, Canada

<u>Supplementary</u>

#### METHODS

All calculations reported in this work involved various methods. For evaluating protein docking for SARS-CoV-2 models, We used Convolutional deep neural network-based approach (Refer Supplementary Data methods). For in vitro screen against Spike S1: ACE2 binding assay. We used ACE2 Inhibition TR-FRET Assay (Refer Supplementary protocol Material and methods Section). For the solvation free energy calculations between different ligand bound complexes we used MMGBSA calculation, and its Mathematical equations implied (is reported in the MMGBSA calculations Supplementary section). For extended Assay conditions, data analysis and molecular descriptors and all other details relevant to our SARS-CoV-2 study have been reported in Supplementary section.

### SUPPLEMENTARY DATA

**Table S1:** Convolutional deep neural network-based approach named DOcking decoy selection with Voxel-based deep neural nEtwork (DOVE)[1] for evaluating protein docking for SARS-CoV-2 models:

| PDB entry | ATOM+GOAP+ITScore* |
|---|---|
| 6WOJ | 0.49124 |
| 6M0J | 0.17618 |

* Input based on atom types, locations, and atoms' GOAP and ITScore energy scores in a 403 A3 cube in the interface area(within 10A), in which the cube center is the interface area center.

**Mutational Landscape:**

Mutational studies have been carried out in an elucidated way by (Li et al., 2020) who analysed 80 natural variants and 26 glycosylation spike mutants using pseudovirus assay. The group concluded glycosylation deletions were less infectious.

**Mac 1 Hydrolytic Activity and Cell culture Experiments:**

Cell culture experiments have revealed that SARS Mac1 is dispensable for viral replication in some cell lines (Fehr et al., 2015, 2016., Erikson et al., 2008, Putics A et al., 2005). In addition to the animal studies reciprocating that Mac1 hydrolytic activity promotes immune evasion and that it is essential for viral replication and pathogenicity in the host (Fehr et al., 2020).

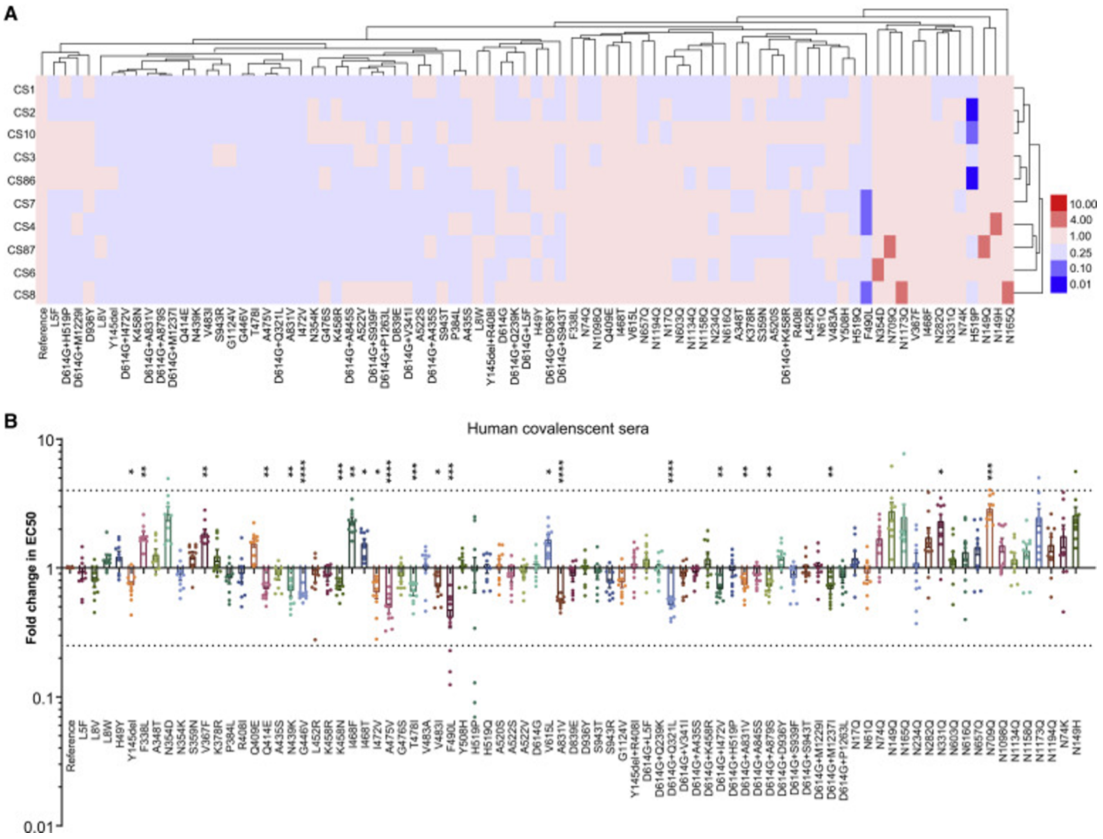


**Figure S1 (A)**. (A) Serial dilutions of 10 patient serum samples were individually mixed with the pseudotyped viruses at 37 C for 1 h before being added to Huh-7 cells for incubation of 24 h to determine the EC50. The experiments were repeated at least 3 times. Hem I software was used to analyzing the data and draw the heatmap. The red and blue boxes indicate the increase or decrease of the neutralization activity as shown in the scale bar.
(B) Summary of the data from ten serum samples, with the values presented as mean ± SEM. The horizontal dashed lines indicate the threshold of 4-fold difference. *p < 0.05, **p < 0.01, ***p < 0.005, ****p < 0.001.

**(Figure S1 (B))**. Human CoVs bind to ADP-ribose with similar affinity where (A and B) depict the binding of human Mdo2 and SARS-CoV, MERS-CoV & SARS-CoV-2 Mac1 proteins

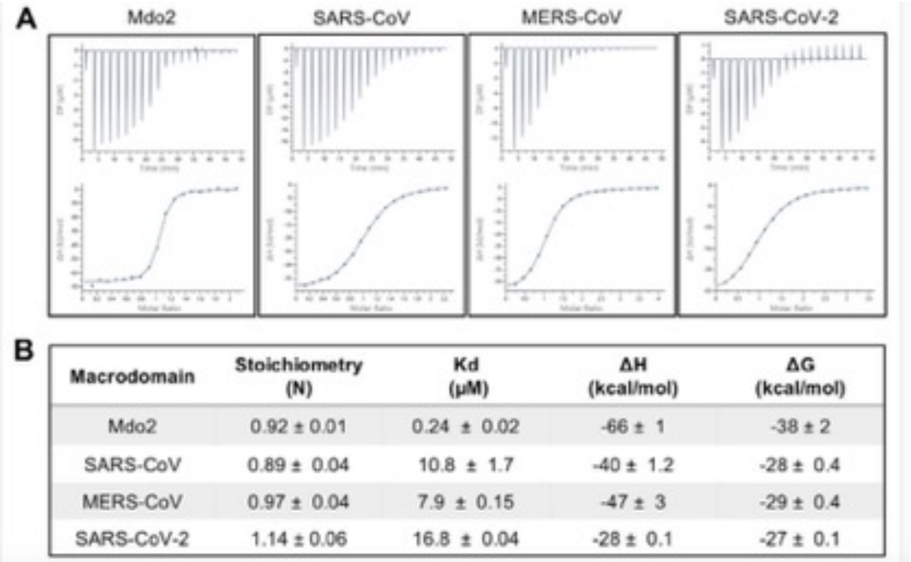


| Macrodomain | Stoichiometry (N) | Kd (µM) | ΔH (kcal/mol) | ΔG (kcal/mol) |
|---|---|---|---|---|
| Mdo2 | 0.92 ± 0.01 | 0.24 ± 0.02 | -66 ± 1 | -38 ± 2 |
| SARS-CoV | 0.89 ± 0.04 | 10.8 ± 1.7 | -40 ± 1.2 | -28 ± 0.4 |
| MERS-CoV | 0.97 ± 0.04 | 7.9 ± 0.15 | -47 ± 3 | -29 ± 0.4 |
| SARS-CoV-2 | 1.14 ± 0.06 | 16.8 ± 0.04 | -28 ± 0.1 | -27 ± 0.1 |

**Figure S2**: Representative Figure., Alhammad YMO, Kashipathy MM, Roy A, Gagné J-P, McDonald P, Gao P, Nonfoux L, Battaile KP, Johnson DK, Holmstrom ED, Poirier GG, Lovell S, Fehr AR. 2021. The SARS-CoV-2 conserved macrodomain is a mono-ADP-ribosylhydrolase. J Virol 95:e01969-20. https://doi.org/10.1128/JVI.01969-20.

**Table S2: List of Compounds and their test range used in this study for WT Spike: ACE2 Binding**

| Compound I.D. | Compound Supplied | Stock Concentration | Dissolving Solvent | Test Range (µM) | Intermediate Dilution |
|---|---|---|---|---|---|
| F5084-0852 | Solid | 10 mM | DMSO | 30 | 4 % DMSO in Assay Buffer |
| F1877-0839 | Solid | 10 mM | DMSO | 30 | 4 % DMSO in Assay Buffer |
| F2619-0022 | Solid | 10 mM | DMSO | 30 | 4 % DMSO in Assay Buffer |
| F1877-1292 | Solid | 10 mM | DMSO | 30 | 4 % DMSO in Assay Buffer |
| F0466-0005 | Solid | 10 mM | DMSO | 30 | 4 % DMSO in Assay Buffer |
| F2085-0027 | Solid | 10 mM | DMSO | 30 | 4 % DMSO in Assay Buffer |
| F0772-2453 | Solid | 10 mM | DMSO | 30 | 4 % DMSO in Assay Buffer |
| F0470-0003 | Solid | 10 mM | DMSO | 30 | 4 % DMSO in Assay Buffer |
| F0827-0193 | Solid | 10 mM | DMSO | 30 | 4 % DMSO in Assay Buffer |
| F2173-1125 | Solid | 10 mM | DMSO | 30 | 4 % DMSO in Assay Buffer |
| Anti-Spike* | Solution | 9 µM | PBS | 0.0001, 0.001, 0.01 | Assay Buffer |

* Reference inhibitor

**TR-FRET assay**

FRET assays utilize donor fluorophores with longer emission times (1-2 ms), such as lanthanide ion complexes, to reduce potential interferences caused by the excitation energy. The optimal distances between donor and acceptor fluorophores are similar to FRET pairs. As TR-FRET assays utilize a time delay between fluorophore excitation and signal acquisition (approximately 50-150 μs in duration) and this acquisition delay is sufficient for avoiding interference by the usual short-lived fluorescence from test compounds.

**Experimental Conditions**

**Table S3:** Enzymes and Substrates used

| Enzyme | Catalog # | Protein lot # | Assay concentration |
|---|---|---|---|
| ACE2-Eu | 100705 | 210701 | 5 ng/rxn |
| WT Spike S1-biotin | 100720 | 200423 | 100 ng/rxn |

**Supplementary Protocol-Materials and Methods**

**Materials**

ACE2, His-Tag, Eu-labeled (BPS Bioscience, #100705)

SARS-CoV-2 Spike S1 Protein, biotin-labeled, (BPS Bioscience, #100720)

3X ACE2: Spike TR-FRET Buffer (BPS Bioscience, #79963)

Dye-labeled acceptor (BPS Bioscience)

**Assay Conditions**

5 µl of ACE2-Eu was incubated with 5 µl of dye-labeled acceptor and 5 µl of inhibitor (see table 2.2) Binding reaction was started by the addition of 5 µl of Spike S1-Biotin, as described in the protocol kit #78281 (Spike S1 [Wild-Type] [SARS-CoV-2]: ACE2 TR-FRET Assay Kit). The reactions were incubated for 1 hour at room temperature, and the TR-FRET signal was finally measured in an Infinite M1000 microplate reader (Tecan) at excitation of 340 nm and emissions at 620 nm and 665 nm.

**Data Analysis**

Binding assays were performed in duplicate at each concentration. The TR-FRET data were analyzed using Prism (GraphPad). Percent inhibition was determined by normalizing the data to signal from negative control wells (Blank, wells without biotinylated ligand, set as 100% inhibition) and positive control wells (No compound, wells in the absence of any inhibitor but with respective buffer, set as 0% inhibition). Data for a reference inhibitor, Anti-Spike, was included as a control for inhibition. The values of percentage activity were plotted on a bar graph.

**Molecular Dynamics Simulation Methods**

To understand the dynamic behavior of the docked complex of the macrodomain and the top leads, Molecular dynamics (MD) simulations of 200 ns (nanoseconds) run for each complex were carried out using the Desmond version 4.4 (Schrodinger suite). Protein was set in water for solvation using the TIP3P box. Following the solvation, charge neutralization was carried out by the addition of $Na+$ and $Cl^-$ ions. Minimization was carried out for the total box which contained protein, water, and ions for 2000 steps using the steepest descent algorithm and then for 3000 steps using the conjugate gradient. The structure was analyzed for its stability and potential energy after the completion of the steps. The total system was then equilibrated. The simulation was performed under the conditions of constant temperature and pressure as the protocol for simulations as defined by (Iqbal et al., 2021) was followed. Briefly, The Martyna-Tobias-Klein scheme was used for pressure coupling. PME algorithm (Case et al., 2008; Darden, York, & Pedersen, 1993) was used for calculating the electrostatic forces; all runs have been performed at 300K at constant volume and temperature (NPT ensemble) under certain periodic boundary conditions.

**Conformational Dynamics**

RMSD of the protein Ca-backbone from its starting position increased to 1.4 A˚for the first 100ns and then became stable around 1.8 A˚in last 100 ns course of the simulation. Deviations got converged during 75 ns simulation time of indicating 0.4 A˚deviation, whereas RMSD of protein-ligand(F0470-0003) complex, it was quite different as we saw the complex had gone conformational change and simulations hadn't equilibrated much during initial 50 ns simulation, however with simulation progression run, as represented in Figure 8(B), structural deviation with a magnitude of >1.8 A˚was visible.

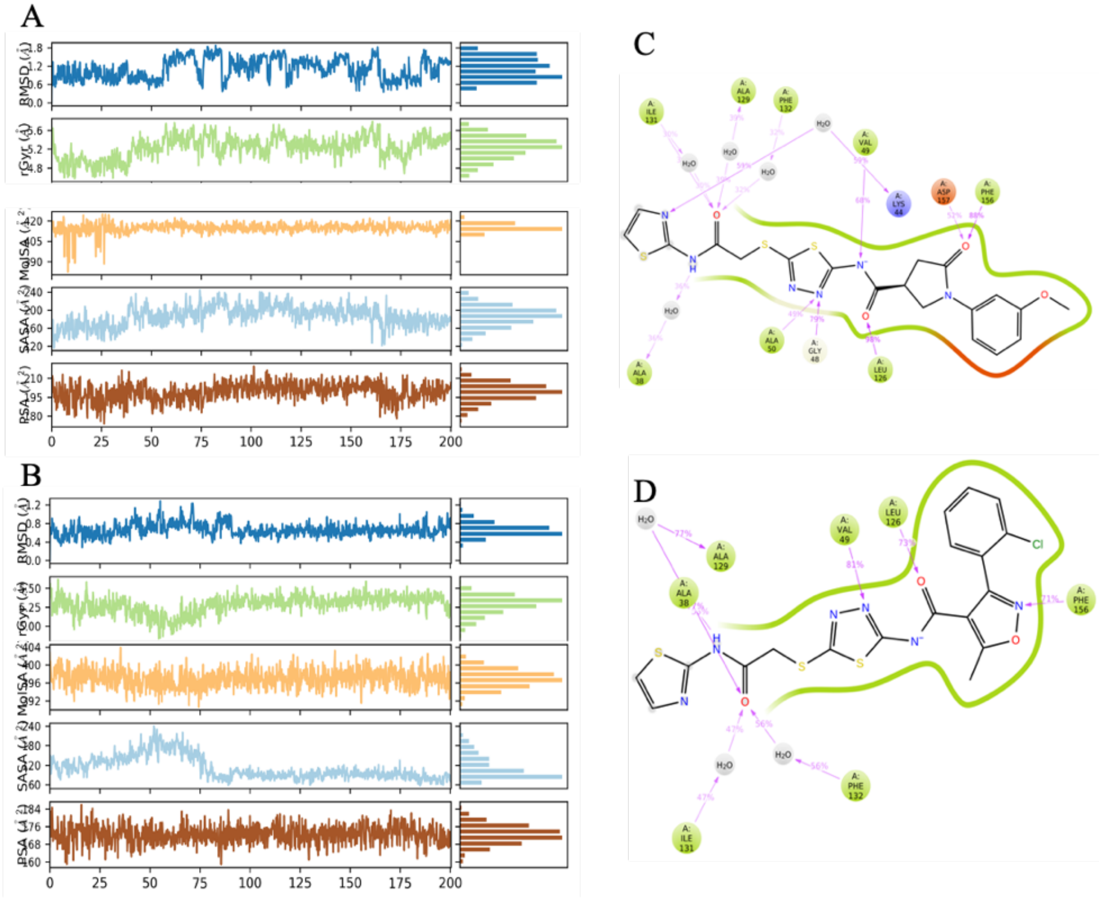


**Figure S3:** (A&B) Ligand properties in terms of PSA, SASA, and rGYR illustrating stability of simulated docked complexes of F1877-0839 and F0470-0003 respectively. (C&D): Simulated Interaction diagram of F1877-0839 and F0470-0003.

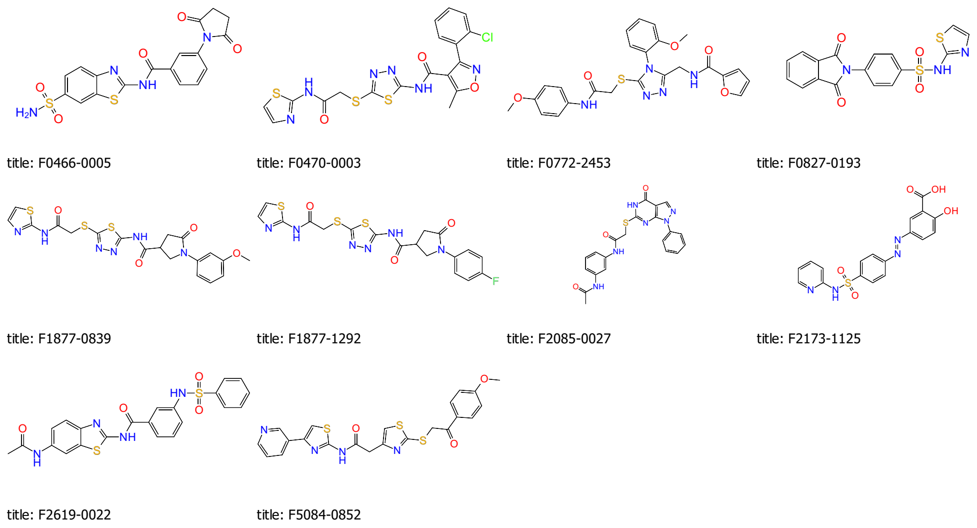


**Figure S4:** Top most selected ligands used for checking the efficacy of Effects of Compounds on WT Spike: ACE2 Binding

All the compounds for HTVS shown in **Figure S4**. were prepared and energy minimized using the Ligprep module of the Schrodinger (Schrodinger 14-2; Sastry, Adzhigirey, Day, Annabhi- moju, & Sherman, 2013), where (with) probable tautomeric and ionization states at pH = 7 ± 1 followed by minimization with OPLS 2005 force field. The ligands shown in Figure S3 were evaluated for checking the efficacy of Effects of Compounds on WT Spike: ACE2 Binding.

**Table S4:** Induced fit Docking results of all the selected compounds and cocrystal at the active site of the macrodomain.

| Compound ID | **Docking Score** | **Glide Energy kcal/mol** |
|---|---|---|
| F5084-0852 | -9.12 | -59.35 |
| F1877-0839 | -12.87 | -78.24 |
| F2619-0022 | -8.25 | -61.34 |
| F1877-1292 | -8.68 | -52.48 |
| F0466-0005 | -9.01 | -60.62 |
| F2085-0027 | -10.32 | -68.03 |
| F0772-2453 | -11.69 | -70.04 |
| F0470-0003 | -9.61 | -75.64 |
| F0827-0193 | -9.08 | -54.86 |
| F2173-1125 | -8.88 | -59.42 |
| Cocrystal | -11.19 | -75.02 |

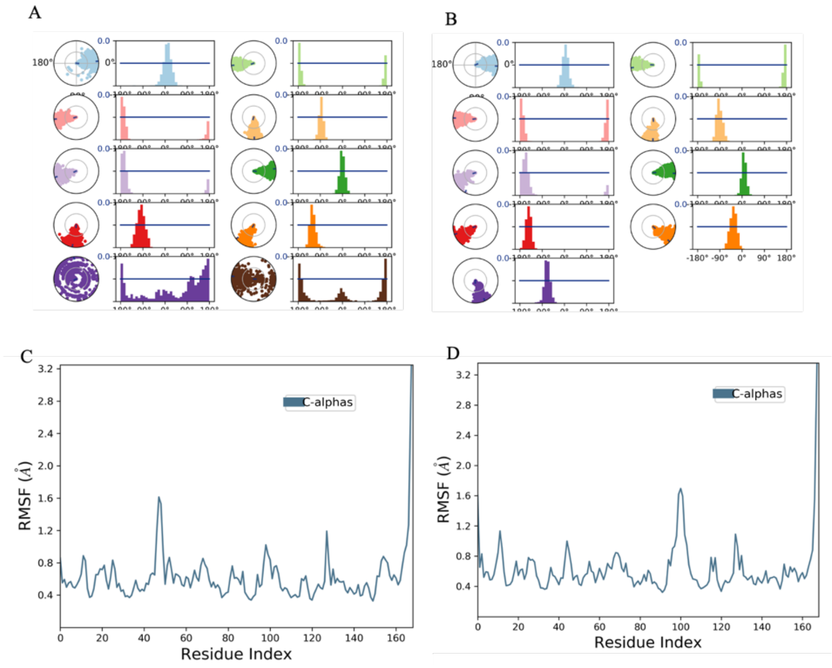


**Figure S5**: (A&B) Torsional profile of the drugs; F1877-0839 and F0470-0003. (C &D): correspond to the respective RMSF fluctuation of the drugs at the active site of the ligand.

#### Table S5: ADMET analysis of the inhibitors used in this study

| Title | FORMULA | MOL_WEI | donor | accptH | QPlogPoc | QPlogPw | QPlogPo | QPlogS | QPlogBB | QPlogK | Human | Perce | RuleOf | RuleOfThr |
|---|---|---|---|---|---|---|---|---|---|---|---|---|---|---|
| F0466-0005 | C18H14N4O5S2 | 430.46 | 3 | 11.5 | 26.468 | 19.954 | 0.352 | -4.669 | -2.793 | -5.76 | 2 | 50 | 0 | 1 |
| F0470-0003 | C18H13ClN6O3S3 | 492.98 | 2 | 10 | 25.24 | 16.359 | 3.262 | -6.744 | -1.492 | -3.34 | 1 | 86 | 0 | 1 |
| F0772-2453 | C24H23N5O5S | 493.53 | 2 | 9 | 24.336 | 14.841 | 4.153 | -5.528 | -1.074 | -1.33 | 3 | 100 | 0 | 0 |
| F0827-0193 | C17H11N3O4S2 | 385.42 | 0 | 7 | 17.522 | 11.43 | 2.399 | -4.276 | -1.347 | -3.22 | 3 | 81 | 0 | 0 |
| F1877-0839 | C19H18N6O4S3 | 490.58 | 2 | 12.25 | 27.589 | 18.377 | 2.323 | -6.227 | -2.026 | -3.74 | 1 | 77 | 0 | 1 |
| F1877-1292 | C18H15FN6O3S3 | 478.54 | 2 | 11.5 | 27.025 | 17.863 | 2.502 | -6.409 | -1.753 | -3.67 | 1 | 79 | 0 | 1 |
| F2085-0027 | C21H18N6O3S | 434.47 | 3 | 9.5 | 26.614 | 18.355 | 2.483 | -5.973 | -2.077 | -3.57 | 3 | 77 | 0 | 1 |
| F2173-1125 | C18H14N4O5S | 398.39 | 1 | 8.25 | 20.914 | 14.074 | 2.456 | -4.564 | -2.559 | -3.89 | 2 | 60 | 0 | 1 |
| F2619-0022 | C22H18N4O4S2 | 466.53 | 3 | 11 | 27.972 | 20.002 | 2.441 | -5.946 | -1.954 | -2.86 | 3 | 80 | 0 | 1 |
| F5084-0852 | C22H18N4O3S3 | 482.6 | 1 | 9.75 | 21.239 | 13.048 | 3.325 | -3.462 | -0.364 | -1.5 | 3 | 100 | 0 | 1 |

**MMGBSA calculations**

MMGBSA method (Miller et al., 2012) was employed to calculate the free energy and energy decomposition per residue, and the $\Delta G_{bind,solv}$ was calculated. For calculating the ligand binding energies and ligand strain energies for docked complexes of leads we used the Prime module of Schrodinger, and was set to (i) Variable-dielectric generalized Born Model (VGSB) which uses Water as continuum solvation model for refinement, (ii OPLS3 - force field, and (iii) minimization sampling method which minimizes all-atom in each residue. The energy of the interaction was calculated from the snapshots taken from the simulation run. Solvation free energy and $\Delta G_{vacuum}$ were also calculated and compared between the different ligand-bound complexes. For the free energy calculation, the following equations given below are used:

$$\Delta G_{binding} = G_{complex} - G_{receptor} - G_{ligand}$$

$$\Delta G_{Tot} = \Delta G_{gas} + \Delta G_{solv}$$

$$\Delta G_{gas} = \Delta E_{ele} + \Delta E_{vdw}$$

$$\Delta G_{solv} = \Delta E_{polar\text{-}solvation} + \Delta E_{non\text{-}polar}$$

Implying… $\Delta G_{bind} = \Delta E_{MM} + \Delta G_{SOL} + \Delta G_{SA}$

where $\Delta E_{MM}$ is the difference in the minimized energies between the mac domain-inhibitor complex and the sum of the energies of the unbound mac domain and inhibitor. $\Delta G_{SOL}$ is the difference in the GBSA solvation energy of the protein–inhibitor complex and the sum of the solvation energies for the unbound mac and inhibitor. $\Delta G_{SA}$ is the difference in surface area energies for the complex and the sum of the surface area energies for the unbound mac domain and inhibitor.

**Binding Affinity Analysis**

The energy of the interaction was calculated from the snapshots taken from the simulation run. Solvation free energy and $\Delta G_{vacuum}$ were also calculated and compared between the different ligand-bound complexes.

**Catalytic Mechanism**

X-ray crystallography augments structure based drug design, and depends upon the accurate model of the shape, electrostatic potential, solvation and flexibility of the target site. Studies decipher that the terminal ribose of ADPr adopts the β epimer, with the C1″ hydroxyl hydrogen-bonded to the backbone nitrogen of Gly48. The proximal ribose ring is stabilized in the pocket by hydrophobic interactions with Phe132 and Ile131, as well as a set of hydrogen bonds with Gly46 (OH2, and Gly49 (OH1). All of these residues are conserved. The distal ribose ring is stabilized by the residues Pro125, Leu126 and Leu127 via hydrophobic interactions. The diphosphate moiety binds between two loops, that cover three segments with high sequence conservation, including a glycine-rich segment (Gly46-Gly47-Gly48) within the former loop, it is conceivable that there is a conformational change in the binding pocket. Structural differences include the peptide flips of Gly48 and Ala129 that allow two new hydrogen bonds with the diphosphate portion of ADPr and a coupled conformational change in the Phe132 and Asn99 side chains to accommodate the terminal ribose of ADPr. Screening of the top most inhibitors at the catalytic active site of macrodomain revealed that the protonation states of catalytic site residues and active site hydrogen bond networks in the docked complexes revealed the molecular basis of functionality relevant flexibility in the active site loops. The large number of bridging waters identified here provides an unmatched opportunity to systematically examine the thermodynamics of water displacement
